# A tale of two springs: contrasting forest soundscapes during the COVID-19 lockdown (2020) and after the record snowstorm Filomena (2021) from Central Spain

**DOI:** 10.1101/2023.10.02.560514

**Authors:** Rüdiger Ortiz-Álvarez, Carmen Leiva-Dueñas

## Abstract

During the COVID-19 pandemic, humanity temporarily retired from the outdoors. The strict lockdown measures in Spain coincided with the onset of the nesting season of birds, thriving in an unusually quiet environment. Here, we have recorded in forests near San Lorenzo de El Escorial (Central Spain) during the lockdown period in 2020, and the closer to normal spring in 2021. We found strong differences in soundscapes by recording year and location, regardless of the effects of meteorology and human mobility. Species altered their behaviour by increasing their calling intensity during 2021 to cope with higher noise levels, however, acoustic activity was generally less diverse and complex. The difference between years was particularly detrimental for the highest-pitched biophony in 2021. We interpret that an extreme snowfall, Filomena, may have caused a mortality event with lasting effects in the community during the 2021 spring. Since extreme climatic events are likely going to keep happening in the area due to climate change, our data is a useful baseline to guide future conservation efforts, and examine how our activity and climate change are changing the soundscapes of Spanish Mediterranean forests.

## 1. Introduction

Sound has traditionally been an overlooked component of ecosystems. Nonetheless, anything that moves makes a sound or vibration: from animal communities (biophony) to geological features (geophony) and human activities (anthropophony). These three types of dynamic signals altogether in a given area form what is called a *soundscape* (Krause 1993; Pijanowski, Villanueva-Rivera, et al. 2011; Pijanowski, Farina, et al. 2011). Although the balance of these sound sources can change over time, sudden global changes are rare. But during 2020, the COVID-19 pandemic created an unprecedented situation where humanity temporarily retired from the outdoors at a global scale (Rosenbloom and Markard 2020; Saraswat and Saraswat 2020), altering human movements and behaviours, a *quietus* or *anthropopause* with substantial acoustic impacts. For example, anthropogenic noise derived from air and land transportation can be an invisible but quantifiable source of habitat degradation (Iglesias-Merchan et al. 2015; Ortiz-Urbina et al. 2020), and bird communities adapt if possible to avoid increasing levels of noise (Ware et al. 2015). In this current context of degradation of natural soundscapes (Morrison et al. 2021), and given decades of observations (Krause 1999), any recordings taken during 2020 are unique. Indeed, studying animal acoustic responses towards this quasi-experiment allowed the opportunity to measure the recovery of the natural world to changes in human behaviour (Montgomery et al. 2021), thriving in a rare acoustic situation that could resemble pre-industrial times. Indeed, reports have already shown fast recovery of soundscapes in some locations, with increased activity of wildlife (Derryberry et al. 2020). Ultimately, ecological measurements from this *anthropopause* serve as a baseline to compare future dynamics and can be extremely helpful to guide conservation efforts.

Since sound is an ecosystem property, recording soundscapes is a profitable strategy to track broad ecosystem reactions to environmental changes. In fact, in the current context of climate change, it is expected that thermal and moisture changes will modify the natural soundscapes and biophony assemblages (Sueur et al. 2019) because weather variations can significantly alter the range of vocalizations between sonoriferous individuals, among other reasons (Larom et al. 1997). In other words, novel weather dynamics might reduce or increase the number of species calls, with implications in the energy invested in other activities required for survival (Sueur et al. 2019). Hence, examining the acoustic assemblage in a seasonal context with variable weather might give us clues about future animal behaviors, particularly in locations with high meteorological variability. For instance, because of its geographical position, Spain is at high risk of increasing extreme climatic events such as droughts, floods, and, as recently shown, record snowfalls (Rodrigo 2010; Lehtonen et al. 2014; AEMET 2021). Indeed, the current climatic trend is leading to more frequent and unpredictable extreme climatic events (Paprotny et al. 2018), damaging not just habitats but its thriving sonoriferous species (Gordon et al. 2018). In this context, it becomes a necessity to register baseline acoustic measurements that allow tracking behavioural and compositional changes in vulnerable areas to help design actions at the local and regional levels, and while broadening our understanding of soundscape vulnerabilities at a global scale (Morrison et al. 2021).

We present the baseline data registered during the spring of 2020 in two forests located in San Lorenzo de El Escorial (Guadarrama mountains, Central Spain). The quarantine measures in Spain lasted from March 14^th^ until May 2^nd^, implying that air and road traffic was drastically reduced, access to green areas was completely shut down, and fortuitously coinciding with the onset of the nesting season of birds. Considering that animal compositions and acoustic behaviour are affected by habitat and vegetation quality (Shaw et al. 2021), baseline soundscapes differ across locations due to vegetation structure and habitat health (Bormpoudakis et al. 2013; Furumo and Mitchell Aide 2019; Robert et al. 2019). Therefore, we compared the soundscapes in two forests with different vegetal compositions: an allochthonous pine tree forest vs. an autochthonous oak tree forest. To test the utility of the 2020 baseline data, we compared it with the following spring, in the same locations and dates, and added a third location farther from the city center as a control. Although the ranges and variability of human activity and mobility 2021 were closer-to-normal, the spring of 2021 came right after the record snowfall ‘Filomena’, an extreme climatic event occurring during January of 2021. Such an event might have caused lasting effects during the subsequent spring when, in addition, the surviving animal community had to thrive with increased levels of noise and human presence. Specifically, to check how different the spring soundscapes from 2020 and 2021 were, we first decomposed soundscapes into multivariate environments, relying on the available toolkit of bioacoustics indexes, spectral and calling compositions. In addition, we developed and included a methodological innovation using a network-based approach that can complement current metrics. We finally determined the extent to which the location, recording year, human mobility and meteorological factors, drive differences in the multivariate environment by using generalised additive models (GAMs). Our starting hypothesis is that the unusual spring of 2020 was particularly favourable for vocal activity, which would be supported by higher acoustic diversities and soundscape complexity, while we predict that the spring of 2021 could be more compromised.

## 2. Materials and methods

### 2.1. Sampling locations and ecosystems

San Lorenzo de El Escorial is a World Heritage Site (UNESCO) and a medium-size town growing around a monument: the Monastery-Palace-Pantheon built by the King Philip II of Spain in the XVI century. The landscape is partially protected under several conservation figures (i.e., *Patrimonio Nacional*), although the pressure of recreational activities in the green areas nearby has been increasing during the last 20 years. Here, the natural ecosystems are characterised by a deciduous autochthonous forest dominated by *Quercus pyrenaica* (‘Herrería forest’, HERR) (**40°34’48.3"N 4°08’56.1"W, 960 m.a.s.l**), and also by an allochthonous pine forest mixed with a few deciduous trees replanted by the end of the XIX century (Miguel del Campo Park, PMC) (**40°35’49.7"N 4°09’32.8"W, 1171 m.a.s.l**). Both locations are easy to access green areas, hence are often used for recreational purposes. Despite this situation, both forests contain diverse yet elusive fauna. In addition, Miguel del Campo Park harbors a small stream with variable water flow. A third location was recorded only during 2021, an allochthonous pine tree forest (close to ‘Cuelgamuros valley’, CMU) (**40°38’06.4"N 4°07’19.0"W, 1193 m.a.s.l**), farther from typical hiking trails and, at a 1.69 km distance from the closest road (**Figure S1**)., as a control, although it was not included in the full analytical workflow. Standard surveys of avian fauna in the areas back in 2003, showed that Herrería forest harboured an outstanding avian density (23.8 individuals/ha), more than the pine tree forests (4.6 and 4.7 individuals/ha) (Moreno-Opo and Seoane 2004). That same survey revealed about 77 bird species thriving in the areas. Such density and diversity are represented mainly by passerines with variable body sizes, such as *Cyanistes caeruleus*, *Parus major*, *Erithacus rubecula*, *Turdus merula*, *Sitta europaea*, *Certhia brachydactyla*, *Troglodytes troglodytes*, *Regulus ignicapilla*, or *Dendrocopos major*, among many others. Despite the constrained conditions of 2020 impeding the logistics of deploying spatial replicates, we argue that a precision temporal analysis, similar to other scientific disciplines such as paleoecology (Leiva-Dueñas et al. 2020), microbial ecology (Felip et al. 1999), or even precision medicine (Kyrochristos and Roukos 2019), can be a powerful strategy to diagnose the processes occurring in our studied locations.

### 2.2. Seasonal, meteorological, and mobility metadata

Daily meteorological data (average temperature, cumulative precipitation, and wind speed) of the nearest Spanish Meteorological Agency (AEMET) station (Robledo de Chavela) to our study was downloaded from R using the package ‘*climaemet’* (Pizarro et al. 2021). The recorded time periods of both years had similar temperature ranges (2020: 10.4 (4.7-15.4° C) vs. 2021: 10.7 (11.1-17°C)) and wind speeds (2020: 1.93 (1.7-3.9) Km/h vs. 2021: 2.1 (1.9-4.2) Km/h), but the spring of 2021 was slightly drier (2020: 2.81 (0.3-27.3 mm) vs. 2021: 1.18 (0-10.7) mm). Regional human mobility was used as a quantification of the anthropogenic activity for each day (**Figure S2**). Data was extracted from ‘Apple mobility reports’ (www.apple.com/covid19/mobility), which uses the relative change of directions requested in Apple Maps for the Madrid region (*Comunidad de Madrid*), compared to the value of January 13^th^ of 2020. The database lacks resolution at the local scale; hence, we consider this metric a proxy of mobility and associated road traffic noise rather than an accurate measurement in each recording location.

### 2.3. Recording scheme and sound pressure analysis

We set automatic recording devices (SWIFT units, from Cornell University) to record mono *wav* files with a sampling frequency of 48 kHz and 16 bits, from a height of ∼1.5 m above ground). The recording scheme was programmed as follows: during 2020, recordings in HERR and PMC started on March 19^th^ and ended on April 12^th^ (a total of 25 days. In HERR, it was extended to include the days between April 17th and April 29^th^, to a total of 36 days). In 2021 the timespan covered the days between March 1^st^ to April 30^th^ (a total of 61 days). The recording schedule was set to continuous at 11:00-12:00 and 18:00-20:00, splitting the recordings into 10-minute fragments, and focusing on the dusk chorus (astronomical dusk between 19:00 and 20:00, although the position of a mountain range causes dusk earlier (R.O-A personal observation). Recordings keep the GMT+1 time zone during the full timeseries. We quantified sound levels in Kaleidoscope PRO v 5.4.6, by retrieving levels along frequency segments (1/3 Octave Bands) and dB values, in each 10-minute fragment. We required adjustments according to calibrated microphone sensitivity (−44dB against a 1kHz 94 dB (=1 Pascal) reference) and gain level settings (+33 dB). Here we report the results with a reference of 0 dB = 20 microPascals. Sound level means and statistics required transforming dB into a linear scale. Statistical differences between years and locations were assessed by using mixed effects models for time-series as implemented in R package ‘nlme’ (Pinheiro et al. 2021). The raw audio dataset supporting the findings of this study is available from the corresponding author upon request.

### 2.4. Spectral profiles

Spectral composition was calculated by first, filtering the signal below 5% with function *afilter* for each 10 minute recording, then the spectrum for each channel was calculated using the *spec* function in ‘*Seewave’* package (Sueur et al. 2008), using a window length of 512, FFT function for faster calculation, returning the Probability Mass Function (PMF), and setting the remaining parameters as default. The 10-minute spectra (a total of 18 fragments) were averaged as replicates with a resolution of two decimals (kHz) to obtain a single representative spectrum per day.

Secondly, for the construction of the daily spectral profiles, we first removed the baseline profile using the function baseline in the R package ‘*baseline’* (Liland et al. 2010), with a modified polynomial fitting of degree 7 and tolerance of 0.01. This fitting was necessary to make comparisons with the CMU time-series because of the increased background noise present in the device (for an example, see **figure S3**).

### 2.5. Estimation of bioacoustic indexes, cluster analysis, and multivariate ordinations

Rather than examining diversity through bird counts, or single bioacoustics indexes, which may not always correlate (Alcocer et al. 2022), we examined the multivariate acoustic environment with three parallel multidimensional ordinations using: (1) bioacoustics indexes and network properties, (2) the spectral profile and, (3) signal-clusters. Firstly, for the estimation of bioacoustic indexes we used the implementation within the Kaleidoscope PRO v. 5.4.2 software to get the following indexes: MEAN, SFM, SH, NDSI, ACI, ADI, AEI, BI, BGN, SNR, ACT; EVN, LFC, MFC, HFC, and CENT (see **Box 1**). To summarise the information of bioacoustic indexes and the two network properties (see full documentation of network analysis in **Supplementary Information S1**), we first standardised and transformed all these using a box-cox transformation and then used a Principal Components Analysis (PCA) with varimax rotation as implemented in the R package *‘psych’* (Revelle 2019). The number of significant principal components (PC) was determined with Parallel Analysis (Horn 1965). Secondly, baseline-corrected spectra were transformed into a relative scale, and then transformed into a Bray-Curtis dissimilarity matrix which was the input of a non-metric multidimensional ordination (Spec.nMDS), both implemented in the R package *vegan* (Dixon 2003). Thirdly, we used call clusters as retrieved by the automatic clustering analysis in Kaleidoscope PRO. We conducted a single analysis under default parameters, detecting signals between 250 to 12000 Hz, and obtaining a table of 902420 signals assigned to 500 clusters. These raw clusters are groups of similar signals, likely coming from a single source or even calls from a single species. However, this method retrieves a high number of false positives (Knight et al. 2017), and specific individual assignments can be wary without manual inspection. Such table does translate directly to the acoustic community, but it is based on probabilities and not in exact numbers, since conducting a manual inspection was not feasible given the high number of signals extracted. In fact, since our aim is not to explain each cluster individually, but to condense all of them into a few community-level metrics, we argue that this automatic analysis can be a valuable option to understand part of the multidimensional acoustic space. To use these automatic clusters as proxies of the acoustic community, we use the three top matches returned by Kaleidoscope PRO (a single signal has assignment probabilities to the three closest clusters). In R, we weight these clusters by the sum of inverse distances of probability assignments of each signal (Cluster ‘abundance’ = ∑*_for each cluster and day_ Max distance - call to cluster distance*). Cluster abundance represents the sum of all the assignment probabilities of signals. Finally, we use the distance-weighted clusters as an input for the nMDS ordination (Kaleido.nMDS).

### 2.6. Generalised additive models to predict the soundscape multivariate environment

Coordinates of individual observations along the multivariate dimensions were extracted from the PCA and the nMDS and linked with their corresponding sites and days to get summarised spatio-temporal information of the soundscapes. Altogether, the 8 summary variables, and the sound levels obtained for frequencies below 2 kHz (caused mostly by geophony and anthropophony) and between 2-8 kHz (where most Biophony occurs), describe the multivariate acoustic environment. To understand how different covariates affect the variability observed in the acoustic time series we constructed generalised additive models (GAMs), as implemented in the ‘*mgcv’* package (Wood et al. 2016). These were used to model the effects of meteorological factors, human mobility, and study location on the temporal trends of the multivariate acoustic space, hence, we did a total of 10 models. The advantage of using GAMs is the potential to evaluate both linear and highly non-linear relationships between response and explanatory variables while additively calculating the component response (Wood 2011). All the explanatory variables were fitted in the 10 models as continuous variables except for location and year, used as nominal variables (each with two levels).

If skewed, response variables were either square root or log-transformed. A Gaussian error distribution was used, together with low-rank thin plate regression splines, to parametrise the smoothed functions of time (Wood 2003). We used the automatic restricted maximum likelihood smoothness selection to select the optimal model coefficients and smoothness parameters (REML) (Wood et al. 2016). Models were also tested for non-linear correlations among explicative variables, i.e. concurvity between smooth terms was assessed. We used the adjusted R^2^ (R^2^ adj) as an indicator of model performance. We carried out model validation by checking plots of residuals versus fitted values, residuals versus location, response versus fitted values, histograms of residuals, and QQ-plots to verify the underlying assumptions of homogeneity and normality. Most models did not violate the assumptions, except for the model having Spec-NMDS2, Kaleido-NMDS1 and Kaleido-NMDS2, for which models output must be very carefully interpreted. Finally, when the normalised residuals showed significant temporal autocorrelation according to plots of a partial autocorrelation function applied to normalised residuals, we introduced an autocorrelation-moving average correlation structure, determined by the *corARMA* function, from the R package ‘*nlme’* (Pinheiro et al. 2021). A methodological summary is displayed in **Figure 1**.

**Figure 1.**
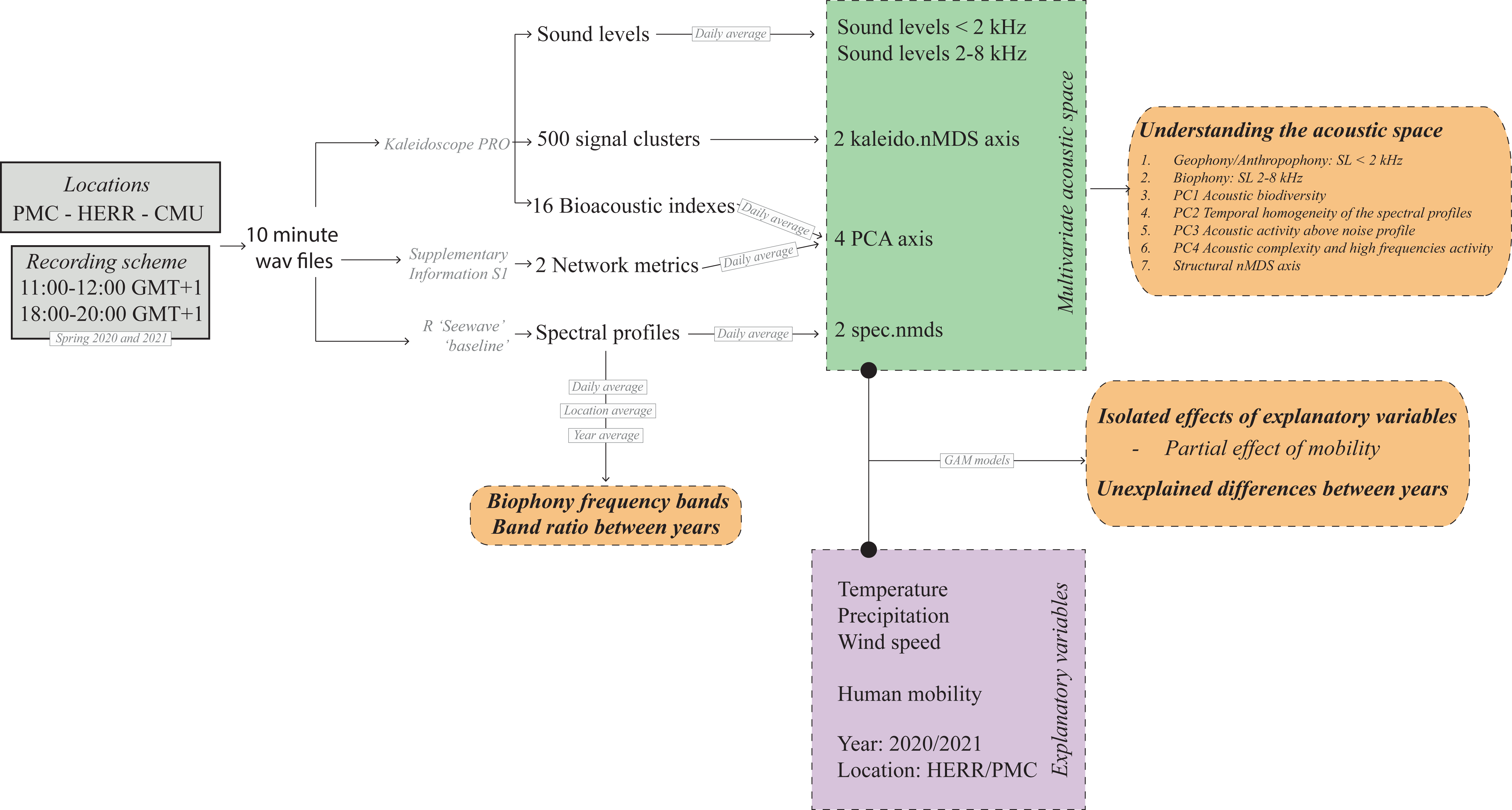
Methodological summary and analytic workflow.

## 3. Results

### 3.1. Time-series overview: spectral dynamics and acoustic space use

Sound pressure shifted between 2020 and 2021 by recording location, and in different frequency ranges (**Figure 2**). Noise levels in frequencies below 1000 Hz (mostly geophony + anthropophony), increased in 2021 compared to 2020, in both locations but more strongly in HERR (**Figure 2B**). According to the GAM results (**Table 1**), regardless of meteorological variability, human mobility contributed significantly to such differences (GAM, p<0.05; **Table 1**). The sound pressure of biophony (2000-12000 Hz) also increased significantly in 2021 (**Figure 2B**) regardless of meteorology and human mobility (GAM, p<0.05; **Table 1**). When comparing the sound pressure between geophony/anthropophony and biophony, we observed that the lower range (20-1000 Hz) was more intense than the biophony. Contrarily, during 2020 in HERR it was the only situation where the biophony had a similar intensity to the geophony/anthropophony (**Figure 2C**).

**Figure 2.**
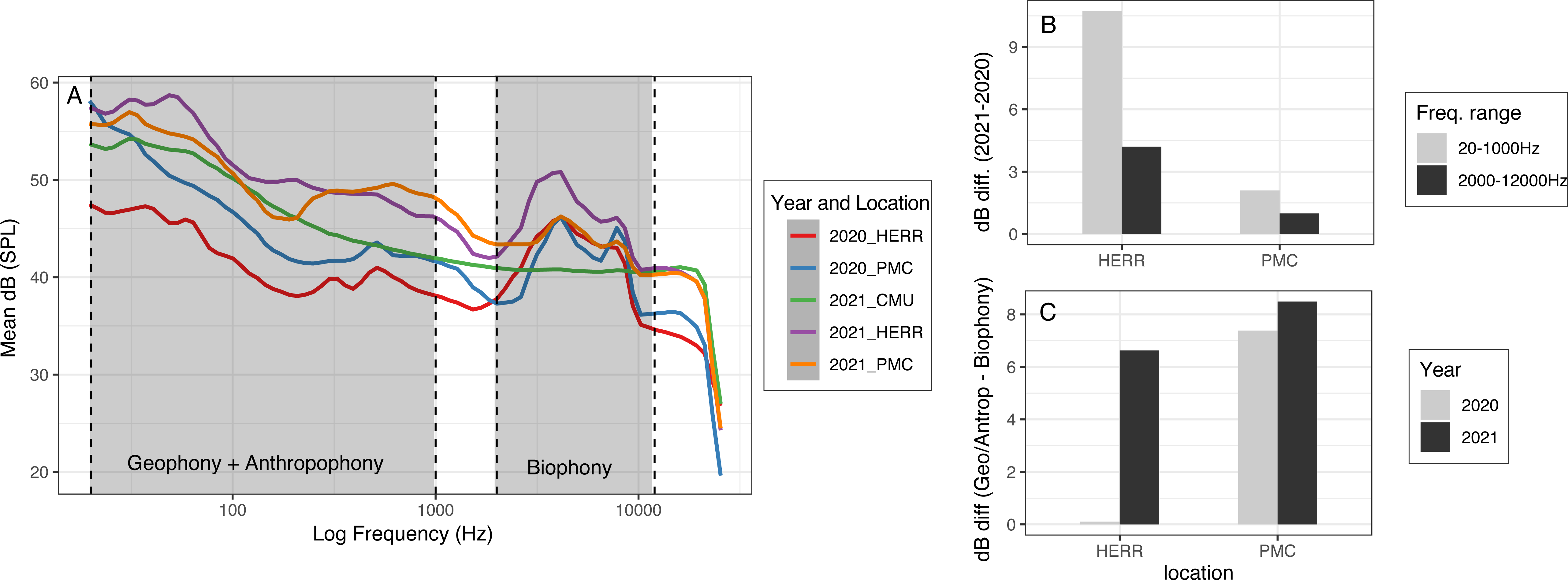
Sound level analysis. (**A**) Mean sound pressure level (dB) across the frequency spectrum (Hz) in each location and year, indicating the frequency ranges that are more likely to include geophony or anthropophony (20-1000 Hz) and biophony (2000-12000). (**B**) Decibel difference between 2021 and 2020 for each location and frequency range. (**C**) Decibel difference between the frequency ranges (20-1000 Hz range minus the 2000-12000 Hz range), for each location and year.

**Table 1.** Summary overview of GAM models, aimed at modelling the multivariate acoustic space. ***: highly significant contribution, ** moderately significant contribution, *significant contribution, .almost significant contribution.

|  | < 2<br>kHz | ~2-8<br>kHz | PC1 | PC2 | PC3 | PC4 | Spec.n<br>MDS1 | Spec.n<br>MDS2 | Kaleido.<br>nMDS1 | Kaleido.<br>nMDS2 |
| --- | --- | --- | --- | --- | --- | --- | --- | --- | --- | --- |
|  | R<sup>2</sup>=<br>0.57 | R<sup>2</sup>=<br>0.39 | R<sup>2</sup>=<br>0.72 | R<sup>2</sup>=<br>0.22 | R<sup>2</sup>=<br>0.73 | R<sup>2</sup>=<br>0.57 | R<sup>2</sup>= 0.45 | R<sup>2</sup>= 0.30 | R<sup>2</sup>= 0.52 | R<sup>2</sup>= 0.55 |
| Mean<br>temperatur<br>e | * | *** | *** |  | *** |  | *** |  | *** | * |
| Mean wind<br>speed | ** |  | *** | * | ** |  | ** |  |  |  |
| Precipitatio<br>n |  | *** | ** |  | * | *** | * |  | *** |  |
| Location |  |  | *** | *** | *** | . |  | *** | *** | *** |
| Mobility | * |  |  | * |  |  | ** |  |  |  |
| Year | . | * |  |  | * | * |  |  |  |  |

To evaluate the biophony of the frequency-amplitude spectrum and isolate it from the noise profile, we conducted polynomic baseline corrections of the daily-averaged spectra, allowing for an equivalent comparison of the biophony regardless of the amount of background noise. After the correction, the main observation is the presence of two dominant biophonic bands, one around 2.5-5 kHz and a second one around 6.5-9 kHz, existing along the whole time-series (**Figure 3A and Figure S4**). These two frequency bands correspond to bird calls (**Figure S5**) of variable peak frequencies, where small birds such as *Regulus ignicapilla* and *Periparus ater* sing within the 6.5-9 kHz band, and slightly bigger birds such as *Turdus philomelos*, *Dendrocopus major* or *Parus major* sing within the 2.5-5 kHz band. Other common birds such as *Cyanistes caeruleus* could contribute to both frequency bands. In addition, there was a lower band at ∼1.1kHz in both locations, which includes voices of people, dog barks, noises coming from car traffic, ambulances or train circulation, or occasional thunders during storms. Such 1.1kHz band was proportionally stronger and more intense in PMC, showcasing the dominant frequency in the water flow from a nearby river. In addition, in PMC, there was a smaller peak at 2.5 kHz present only in 2021.

**Figure 3.**
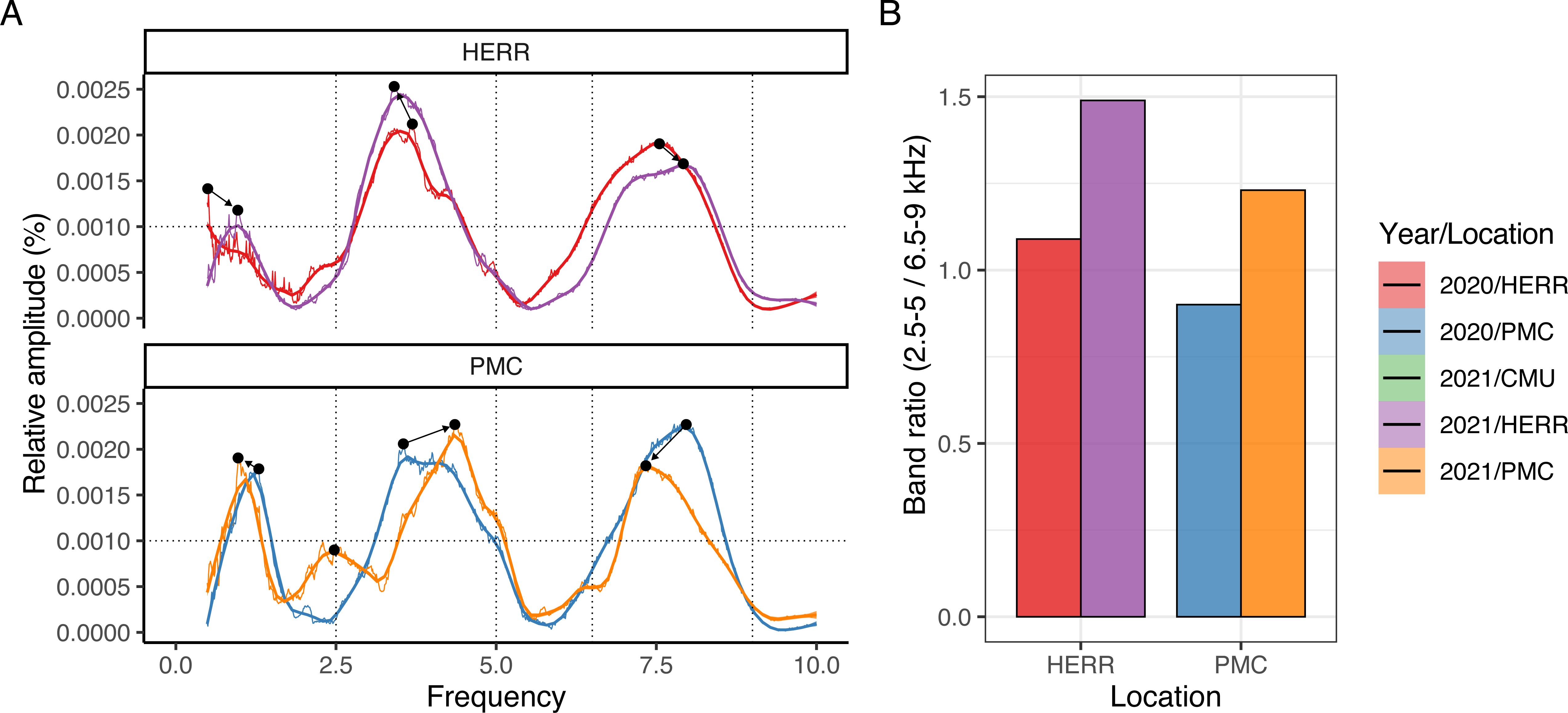
Dynamics of the frequency bands with dominant biophonic activity (Lower: 2-5.5kHz; Higher: 6.5-9kHz). (**A**) Comparison of frequency-amplitude average spectra, highlighting with dotted lines frequency and amplitude thresholds of the dominant frequency bands, and with black dots the maximum amplitude peak. Arrows show the peak switch between years. (**B**) Peak amplitude ratio between the 2-5.5kHz and the 6.5-9kHz for each location.

The most consistent result, regarding the relative amplitude of the biophony bands, is a change on the amplitude ratio in both locations (**Figure 3B**): while in 2020 both bands had similar relative amplitudes, during 2021 the 6.5-9 kHz band was proportionally less intense than the 2.5-5 kHz band. Conversely, there were frequency displacements in the biophony bands when comparing 2020 vs. 2021, but in opposite directions on each location.

### 3.2. Principal components analysis of acoustic metrics

We used the bioacoustic indexes and network properties (designed for this work) to categorise the acoustic space, and summarised them using PCA. Four principal components accounted for 84% of the variation, and showed differences between years and locations (**Figure 4A-B**). Each component is understood as an acoustic trait of the soundscape. For instance, PC1 comprised acoustic diversity measurements (i.e., ADI or AEI) and indicators of acoustic activity of fauna (i.e., NDSI, MFC or BI), displaying a gradient between high acoustic biodiversity (positive loadings) vs. less biodiverse or flatter spectra (negative loadings). In other words, it represents the activity and diversity of the biophony. We analyzed the effect of location and year on the different PC scores through mixed effects models for paired time-series. On the PC1 scores, there was a statistically significant interaction between the effects of location and year (t = 2.94, p < 0.01). In both years, HERR had significantly higher scores than PMC, and that within HERR, year 2020 had significantly higher scores than 2021. We found no differences between years within PMC. The location CMU had the most negative PC1 score values and clearly separated from the other two locations, pointing to less acoustic diversity. According to the GAM model, meteorological variables and the recording location explain most of the variability of PC1 (**Table 1**).

**Figure 4.**
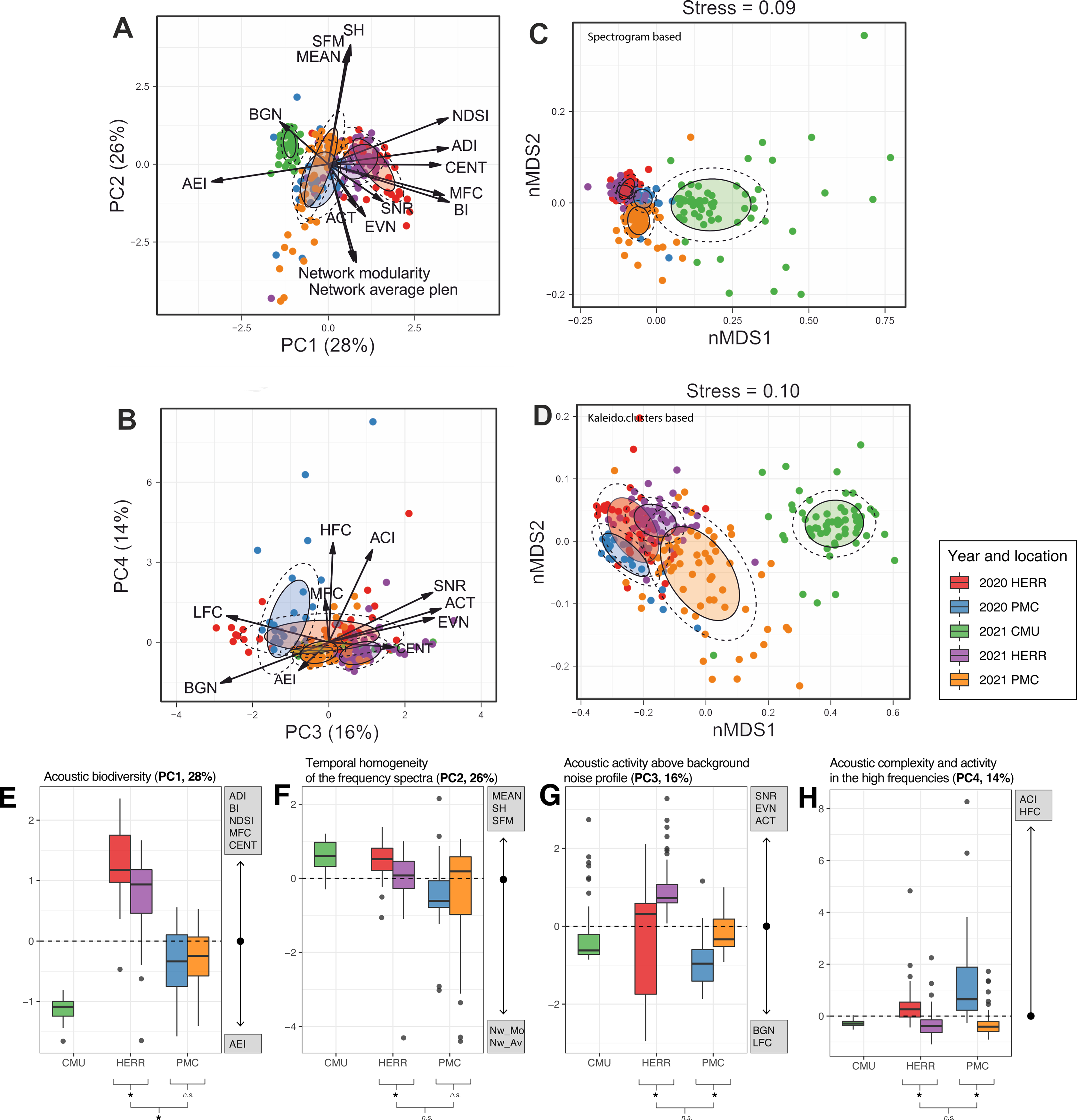
Multivariate ordinations results. (**A-B**) Principal component analysis, (**C**) nMDS based on Kaleidoscope cluster probabilities, (**D**) nMDS based on frequency spectra. (**E-H**) Four PCA loadings (weight > 65%), the contributing variables, (both positively and negatively) axis interpretation, and whether the factors ‘year’ and ‘location’ significantly separated values in each PC axis.

The bioacoustic indexes in PC2 point to a gradient showing soundscape variability at a 10-minute scale. Positive loadings indicate more temporally homogeneous soundscapes with higher mean frequencies and possibly multiple frequency peaks (i.e. higher frequency entropy (SH)). And on the other side, negative values may reflect soundscapes dominated by fewer and specific frequency bands, with a tendency towards lower pitches, and synchronically divided into different events at the 10-minute scale (see **Supplementary Information S1**). The scores from this component were significantly different between locations, and had an interaction with year (t= 2.18, p = 0.032), showing contrasting differences between locations. The highest values were found during 2020 in HERR, alongside CMU, and the lowest during 2020 in PMC. Both HERR and PMC were similar during 2021. The GAM models revealed that human mobility had an effect on the variability of PC2 (**Table 1**).

PC3 has positive loadings for ACT, EVN and SNR, while negative ones for BGN and LFC, reflecting the activity above the noise profile. The mixed-models indicated that PC3 scores had statistically significant effects of location (t = 2.55, p < 0.01) and year (t = 5.50, p < 0.001), with HERR and 2021 having higher PC3 scores values, and hence higher probability of more intense sounds. The location CMU had PC3 score values similar to PMC. The GAM model revealed an effect of the recording year, independent from meteorological variability, and no effects of human mobility on PC3 (**Table 1**).

Finally, PC4 was the only component associated to ACI and HFC. There was a statistically significant difference by year (t= 2.63, p < 0.01) on the PC4 scores. Both locations had lower scores in 2021 than in 2020. In addition, during the year 2020, the location PMC had significantly higher scores than HERR, but there is no evidence to suggest a difference between locations during 2021. The location CMU had PC4 score values that did not differ from PMC and HERR. In most cases, major abrupt peaks correspond to meteorological events (such as intense precipitations or sudden temperature drops) (see **Figure S4**). The GAM model revealed that part of the differences between years were independent from meteorological variability, and no effects of human mobility on PC4 (**Table 1**).

### 3.3. Non-metric multivariate ordinations

The other multidimensional ordinations (**Figure 4C,D**) were constructed using the daily-spectrum profiles after a baseline correction, as displayed in **Figure S4** (Spec.nMDS), and using the approximate call composition by day extracted from the Kaleidoscope PRO cluster analysis (Kaleido.nMDS). Hence, Spec.nMDS reflects part of the patterns and variability shown in **Figure 3D and Figure S4**, while Kaleido.nMDS may reflect changes in call-cluster composition/behavior (**Figure 3C**). Years and locations were well separated using the spectral profiles and an ANOSIM analysis (R_HABITAT_=0.65*, R_HERR_=0.23*, R_PMC_=0.41*, *:p<0.001), and contrary, the call compositions (Kaleido) of HERR were not different between years (R_HABITAT_=0.67*, R_HERR_=0.07, R_PMC_=0.22*, *:p<0.001). According to GAM models, only in Spec.nMDS1, human mobility has a partial effect; while the covariates explaining differences in the 4 axes were meteorology and location (**Table 1**).

## 4. Discussion

Soundscapes are an emergent property of ecosystems, where the biophony represents both the structure of the animal community and its behaviour. Hence, any observed differences in the biophony would indicate either community structural differences, changes in animal behaviour, or both (Pijanowski, Villanueva-Rivera, et al. 2011; Gasc et al. 2013). In theory, since this work takes place in two habitats along the same period with similar weather ranges, we would not expect strong structural community differences between years without the effects of other perturbations. Indeed, the biophony showed two prominent frequency bands, but their dynamics differed according to the recording years and locations (**Figure 5**). Here, soundscapes between 2020 and 2021 had substantial differences in the studied locations. However, given our observations, finding the exact causes of these differences is not trivial.

**Figure 5.**
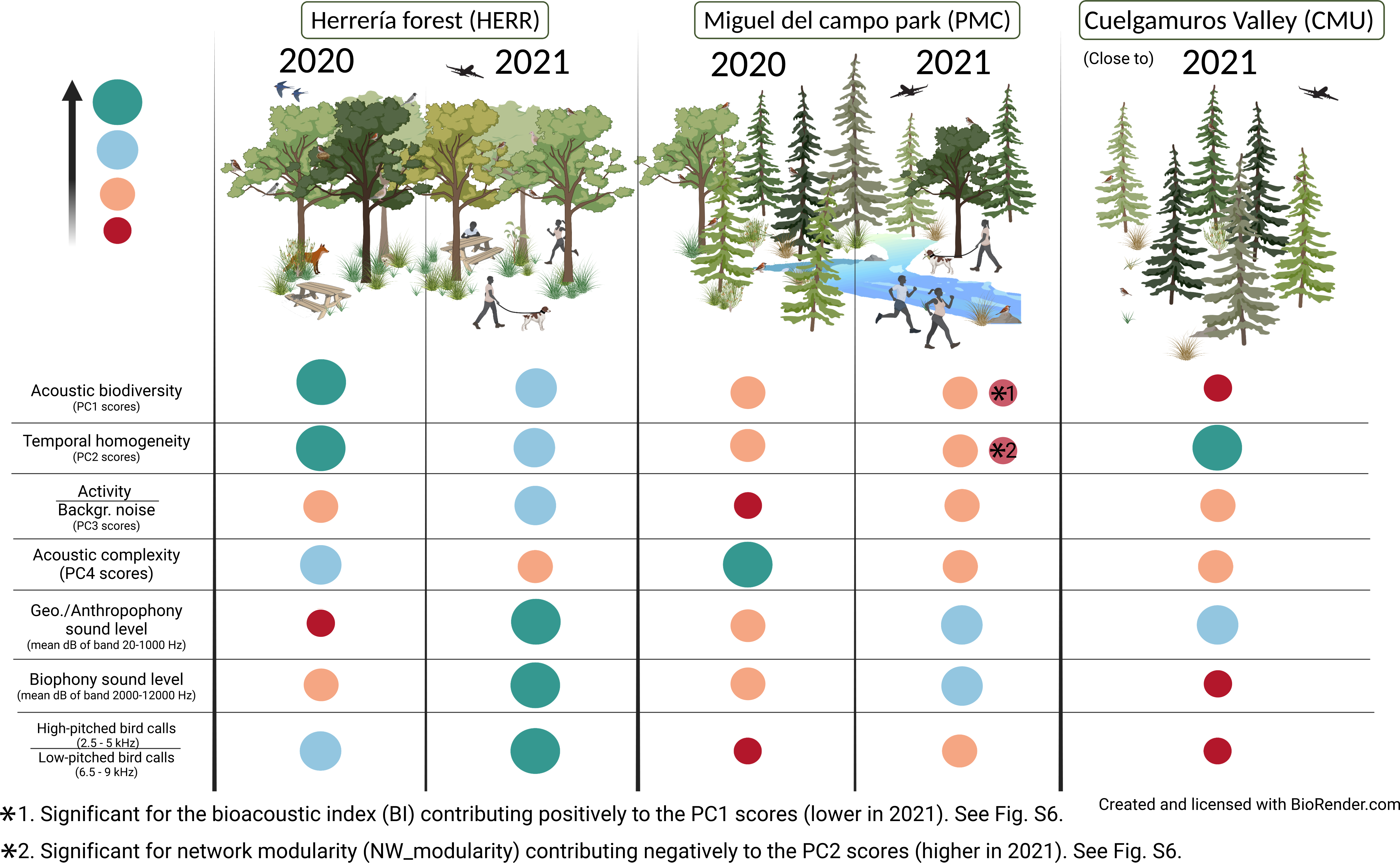
Results summary. Graphical representation of the three habitats (HERR, PMC and CMU) between the two years (2020 vs. 2021), and with the major significant differences observed in the multivariate acoustic space. Ball size does not represent absolute magnitudes between variables, and it is a strict qualitative impression of the main results.

### 4.1. Effect of lockdown and human mobility changes on animal behaviour

The obvious difference between both years was the ‘lockdown’ effect that took place in 2020, when human mobility in the outdoors was almost reduced to zero, causing a decrease in the geophony/anthropophony in 2020, with a maximum difference of ∼10 dB in the *Herrería Forest* compared to the closer-to-normal 2021. The GAM models allowed us to detect a direct effect of human mobility in the frequency range below 2 kHz, and frequency composition, most likely either anthropogenic noises or lower-pitched animal species more prone to suffer acoustic interferences. Also, despite the GAM models show that ‘year’, and not ‘human mobility’ is contributing to explaining the differences between years in PC3 and in the 2-8 kHz sound levels, the expectation would be that sonoriferous species may have adjusted by increasing their calling intensity to avoid masking effects and optimise sound transmission, coping with increasing levels of noise (Brumm 2004). We should note that given the contrasting ranges of human mobility in 2020 vs. 2021, the fact that the quantitative metric is an average of the region and was not locally measured, and that some GAM models have a low R^2^, perhaps differences between years in PC3 and 2-8 kHz sound levels could be due to more local unmeasured lockdown effects, or part of some other unmeasured phenomenon. We conclude that anthropogenic noise is a major source of disturbance reaching protected areas (Barber et al. 2011), affecting the behaviour of sonoriferous fauna needing to invest more energy on communication than under a quieter situation (Pieretti and Farina 2013).

In addition to noise, human presence and conversations can cause mild behavioural changes in animals, temporarily decreasing their calling activity (Karp and Guevara 2011). In our study, we detected an effect thanks to the network metrics developed for this study (**Supplementary Information 1, Figure S6**). In 2021, the moments of highest human mobility caused the more sudden and short stochastic acoustic events (i.e., a plane, passing trains, people talking, or dogs barking) while decreasing the mean frequency of the soundscape fragment (PC2). In other words, human mobility increased the temporal heterogeneity of soundscapes. In addition to human mobility, other factors could contribute to soundscape heterogeneity, such as precipitations or thunder. These geophonic or anthropogenic acoustic events may cause a behavioural reaction and temporal adjustment of calls (Brumm 2004; Nemeth et al. 2013; Ortiz-Urbina et al. 2020), making the soundscape more heterogeneous during the 10-minute segment. Unfortunately, in our study, solely with the network-based metric, it is challenging to separate the intrinsic effect of having more short stochastic acoustic events, which may have indeed lower frequencies, from a pure behavioural response. But theoretically, the network-based metric should be capable of detecting sounds that tend to occur temporally closer to each other, including bird calls. Hence, variability on this metric has the potential to detect short-term behavioural changes: the transition from night to dawn (Gil and Llusia 2020), animal call coordination (Araya-Salas et al. 2017; Araya-Salas et al. 2019), call competition (Farina and Pieretti 2014; Symes et al. 2016; Araya-Salas et al. 2017), or reactions to perturbations (Sethi et al. 2020), adding to the knowledge on soundscape structures and its temporal acoustic assembly (Deichmann et al. 2018; Francomano et al. 2020). We want to highlight the potential of this tool and approach, but it should be further evaluated with more detail and tuning up parameters in the future.

### 4.2. Community structure differences potentially caused by an extreme snowstorm

The acoustic environment’s variation between years cannot be explained solely by meteorological variables, location or human mobility. In 3 GAM models, we observed partial effects by the recording year in the sound levels of biophony (2-8 kHz), PC3 and PC4, suggesting a structural difference in soundscapes beyond the impact of recording location, regional human mobility or season dynamics.

The following argumentation attempts to explain such differences. In both locations, the proportion between the prominent biophony bands showed the same trend in 2021 when compared to 2020: a decline in the relative intensity of the 6.5-9 kHz frequency band. To explain this pattern we need to acknowledge that the 6.5-9 kHz frequency band probably corresponds to songs from smaller birds while the 2.5-5 kHz frequency band corresponds to medium-sized birds, which fits a rough manual exploration through BirdNET (Kahl et al. 2021) (**Figure S5**), and as a global analysis of bird song frequency has demonstrated (Mikula et al. 2021). Considering this, we need to trace back to the previous winter, when the sudden and extreme snowstorm ‘Filomena’ (an event only comparable to a snowfall from 1971 (Pereda and Santos 2021) may have triggered mortality of birds biasing by body size, increasing mortality of smaller birds, and therefore, changing the acoustic structure of soundscapes in the later spring. Indeed, winter weather is responsible for bird mortality through drastic temperature decreases and lower food availability (Jones et al. 2003). Small birds accumulate less fat in regions with mild winters, when the perceived risk of starvation is low. Therefore, a sudden and unpredictable harsh winter could have caused higher mortality in those bids that did not accumulate enough fat for survival (Rogers and Reed 2003); and the repercussions impacting bird breeding productivity could still be observed during the following spring (Saracco et al. 2019).

The extreme snowstorm hypothesis would explain some of the observed differences between years in the GAM models: the decrease in acoustic complexity and higher frequencies as represented in PC4, and also the decline of the biophony in the 6.5-9 kHz frequency band. The extreme snowstorm hypothesis hardly explains why PC3 and the biophony sound levels are higher in 2021 than in 2020, since bird mortality would cause lower acoustic activity and sound levels; hence, we argue that in these two models the difference might be due to local unmeasured effects of lockdown, as already discussed. In contrast, although not shown by the models, perhaps the decrease in acoustic biodiversity (PC1) observed solely in the *Herrería Forest* could be due to such an extreme event, since it reflects the structure of the soundscape (interestingly, the single index BI, contributing to PC1, also decreases significantly in this location in 2021 (**Figure S6**)). Given that extreme climatic events are increasing every year (Lehtonen et al. 2014), we hint that although the acoustic community has some capacity of adaptation, the level of stress could cause structural responses in the community with consequences for the whole ecosystem in the short term (Bellard et al. 2014). Extreme climatic events could be one of the reasons behind the global acoustic diversity loss during the past 25 years, as observed using soundscape reconstructions (Morrison et al. 2021), in addition to the steady increase in temperatures (Sueur et al. 2019).

### 4.3. The effect of habitat type on soundscape structure and dynamics

The biophony is an ecosystem trait reflecting habitat health (Bormpoudakis et al. 2013; Figueira et al. 2015; Gordon et al. 2018). Although the term ‘health’ might be a bit subjective, it is a fact that the biophony reflects the composition and activity of wildlife, which prefers mature (Figueira et al. 2015) and undisturbed habitats (Ware et al. 2015). In our results, we compare three forests, an autochthonous deciduous forest, with two replanted pine tree forests at 200 m higher altitudes, showing that acoustic biodiversity and intensity were higher in the deciduous forest, the *Herrería Forest.* This result agrees with previous studies showing how higher canopy covers and tree heights, summed to forest maturity can positively affect the biophony (Figueira et al. 2015; Hao et al. 2021). This higher ‘health’ of *Herrería Forest* compared to the other locations was consistently observed in 2020 and 2021, but furthermore, this ‘health’ reached the highest values in 2020.

### 4.4. Environmental forces driving soundscape assembly and their implications

In addition to years and locations, which were the strongest differentiators of the soundscapes, we have been able to quantify the impact of meteorological variables, which have an effect in 9 out of 10 GAM models. Spring in Central Spain is the moment of the year when habitats activate their vegetation, and temperature slowly increases. A seasonal succession is aligned with increased acoustic biodiversity, which increases in warmer, drier and less windy conditions, possibly increasing breeding productivity of birds (Saracco et al. 2019). Under such a situation, sonoriferous species form a well-mixed stable chorus, with less heterogeneity and higher pitches, that might be interrupted by windy conditions or anthropogenic noises. Then, fauna might go silent briefly and rearrange, to avoid wasting energy in a non-optimal situation (Popp et al. 1985; Brumm and Slabbekoorn 2005; Brumm 2006; Dooling and Popper 2016). As hinted during all this work, soundscapes are affected by a complex multifactorial environment, and if meteorological changes keep changing due to climate change, sonoriferous species will keep adapting, modifying the whole soundscape and its components (Sueur et al. 2019).

## 5. Conclusions and perspectives

The baseline acoustic data gathered during the COVID-19 lockdown in the Spring of 2020 was useful to track wildlife behaviour under the absence of anthropogenic noise, and observe changes during the following years. Compared with 2021, we observed a marked change in the acoustic space, by increasing sound levels, although the biophony did not increase as much as the anthropophony, and even showed lower acoustic complexity, pointing out to a mortality event caused by the snowstorm Filomena. This event might have affected more strongly higher-pitched birds. Without the 2020 data as a baseline we could not have detected all these changes, and this is only comparing two years. In the future, longer time-series could refine answers at the local-level, in the current global context of declining acoustic biodiversity. We want to stress out that the acoustic monitoring of wildlife, and their reactions to a changing environment, can help implement measures to implement those locations with a higher potential for higher acoustic diversities. Using our data to develop policies and conservation guidelines could make local tourism more sustainable with regards to the acoustic environment, *tourism-of-the-ear*, enhancing the human connection to nature through sound.

## Acknowledgements

This work has been funded by the National Geographic Society (grant NGS-83428R-20). Fieldwork permits for *Herrería Forest* were granted by Patrimonio Nacional. The authors thank A.Llera, B.Vidal, D.K.Ruiz, and X.Pacios and for fieldwork assistance. We specially thank Laurel B. Symes for her support to the project and helpful comments during early stages. The authors declare no conflict of interest.

## BOX 1.

### Essential definitions

**Ecoacoustics** (Sueur and Farina 2015). Umbrella discipline that studies the emergence of environmental sounds as a proxy or marker for ecological processes, covering all ecological organization levels.

**Soundscape ecology** (Pijanowski, Villanueva-Rivera, et al. 2011). The combination of animal signals, geological features, and anthropogenic impacts, non-randomly arrange to form soundscapes. Soundscape ecology focuses on its study with a landscape perspective.

**Frequency spectrum.** Distribution of sound amplitude among a frequency gradient, given in Hertzs (Hz).

**Acoustic diversity index (ADI)** (Villanueva-Rivera et al. 2011). ADI is calculated by dividing the spectrogram into bins and taking the proportion of the signals in each bin above a dB threshold. The ADI is the result of the Shannon index applied to these bins. In this work, we divided the 20-20000Hz spectrogram into 500Hz bins, and used a threshold of −50db.

**Acoustic Evenness Index (AEI)** (Villanueva-Rivera et al. 2011). AEI is calculated by dividing the spectrogram into bins and taking the proportion of the signals in each bin above a threshold. The AEI is the result of the Gini index applied to these bins. In this work, we divided the 20-20000Hz spectrogram into 500Hz bins, and used a threshold of −50db.

**Bioacoustic Index (BI)** (Boelman et al. 2007). Index values are a function of both the sound level and the number of frequency bands used by the avifauna.

**Normalised Difference Soundscape Index (NDSI)** (Kasten et al. 2012). Ratio between common anthropophony and geophony frequencies versus biophony frequencies. In this article, we used a 20-1000 Hz range for the geophony and anthropophony, and a 1000-10000 Hz range for the biophony.

**Acoustic complexity Index (ACI)** (Pieretti et al. 2011). This index is significantly correlated with the number of bird vocalizations. It is cumulative by time, and quantifies song regardless of strong anthropogenic/geophonic noises, accounting for variability in the intensity values.

**Background Noise (BGN)** (Towsey 2017). Background noise estimate of the time domain signal in dB.

**Signal to Noise Ratio (SNR)** (Towsey 2017). The difference between the maximum decibel value in the decibel envelope and the decibel value of BGN.

**Activity (ACT)** (Towsey 2017). The fraction of values in the noise-reduced decibel envelope that exceed the threshold, θ = 3 dB.

**Events per Second (EVN)** (Towsey 2017). A measure of the number of acoustic events per second. An acoustic event is defined as starting when the decibel envelope crosses a threshold, θ, from below to above, where θ = 3 dB.

**Frequency cover** (Towsey 2017). The fraction of noise-reduced spectrogram cells that exceed 3 dB in the low-frequency band **(LFC)** (1-1000 Hz), mid-frequency band **(MFC)** (1000-8000 Hz), and high-frequency band **(HFC)** (8000–11025 Hz).

**Spectral centroid (CENT)** (Towsey 2017). The average amplitude-weighted frequency relative to Nyquist computed individually on each frame ignoring frequencies below 500 Hz.

**Mean frequency (MEAN)** (Sueur et al. 2008). The amplitude-weighted mean frequency.

**Shannon Spectral Entropy (SH)** (Sueur et al. 2008). A noisy signal will tend towards 1, whereas the Shannon spectral entropy of a pure tone signal will tend towards 0.

**Spectral flatness measure (SFM)** (Sueur et al. 2008). The flatness of a frequency spectrum. A noisy signal will tend towards 1, and a pure tone will tend towards 0.

**Time-adjacency network (see Supplementary Information S1).** Network graph constructed by how the frequency spectra of time segments (1 second) are adjacent to others during a longer period (10 minutes).

**Network modularity (NM)** (Pons and Latapy 2005). Modularity of a network measures how good is the network separated into node groups. In this article, nodes are clusters of similar time segments (1 second), and a strong separation (a high modularity value) indicates that a soundscape (10 minutes) has acoustic events that divide it or structure it. In other words, the soundscape is more heterogeneous.

**Network average path length (NAPL)** (Csardi 2008). Calculation of the shortest paths in the network between time segments, averaging the shortest paths for each time segment. A network with higher path lengths, is a network where time-segments are more separated, hence, the soundscape is more heterogeneous and has differences at the given time resolution (i.e., 10 minutes).

## Notes

### Competing Interest Statement

The authors have declared no competing interest.

