## Supplementary figures and images for "A tale of two springs: contrasting forest soundscapes during the COVID-19 lockdown (2020) and after the record snowstorm Filomena (2021) from Central Spain"

### Figure S1

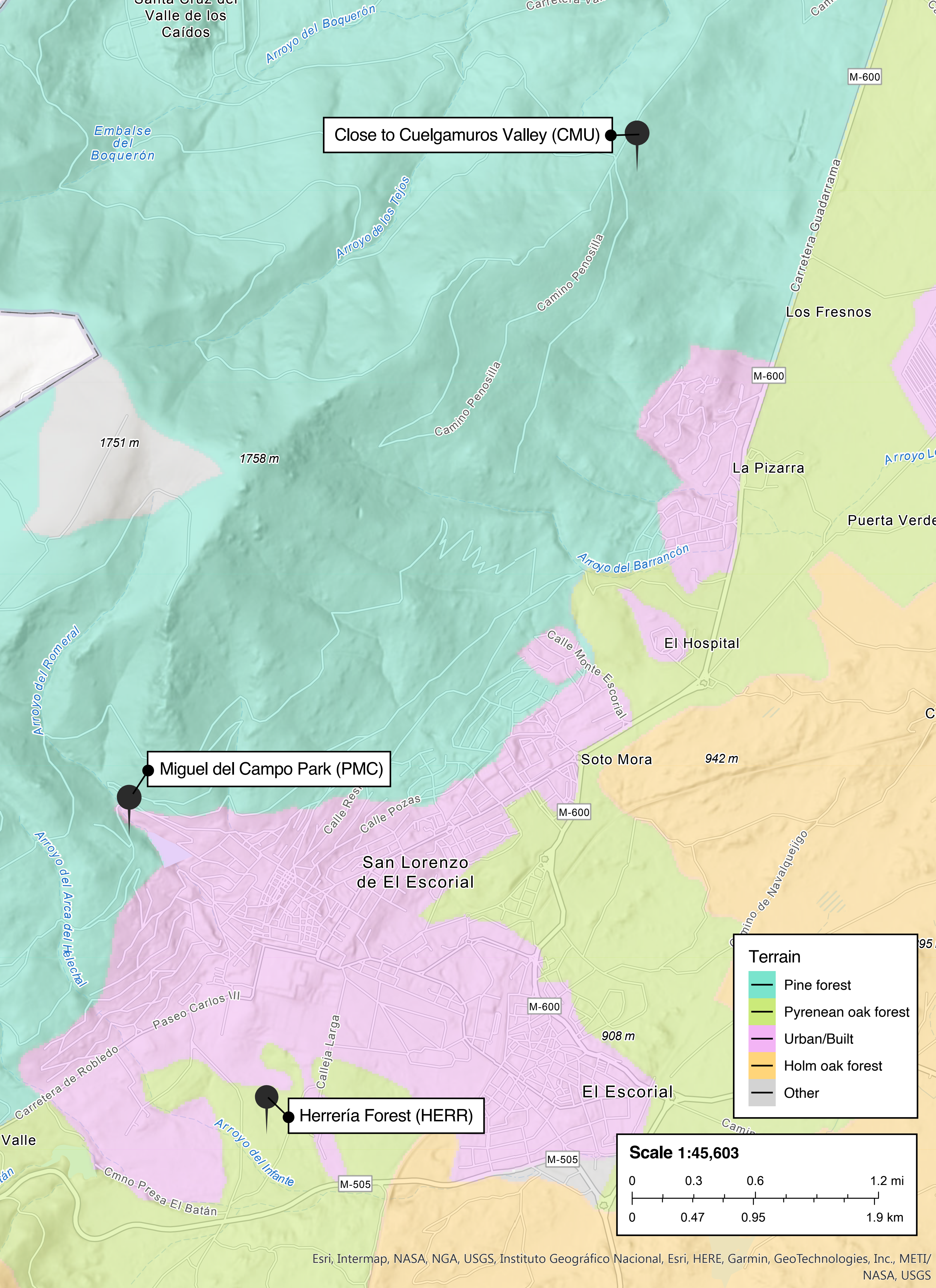

### Figure S2

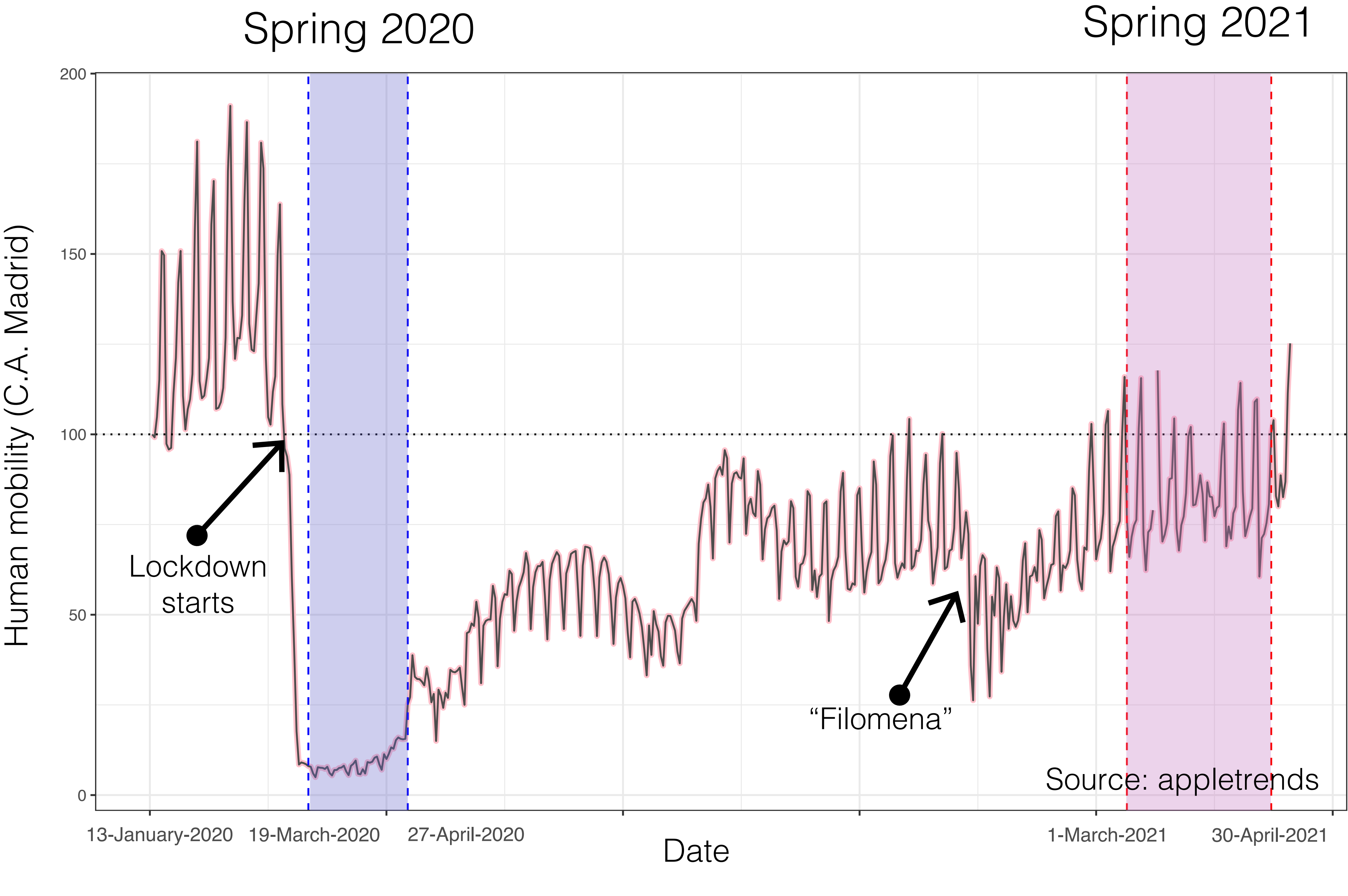

### Figure S3

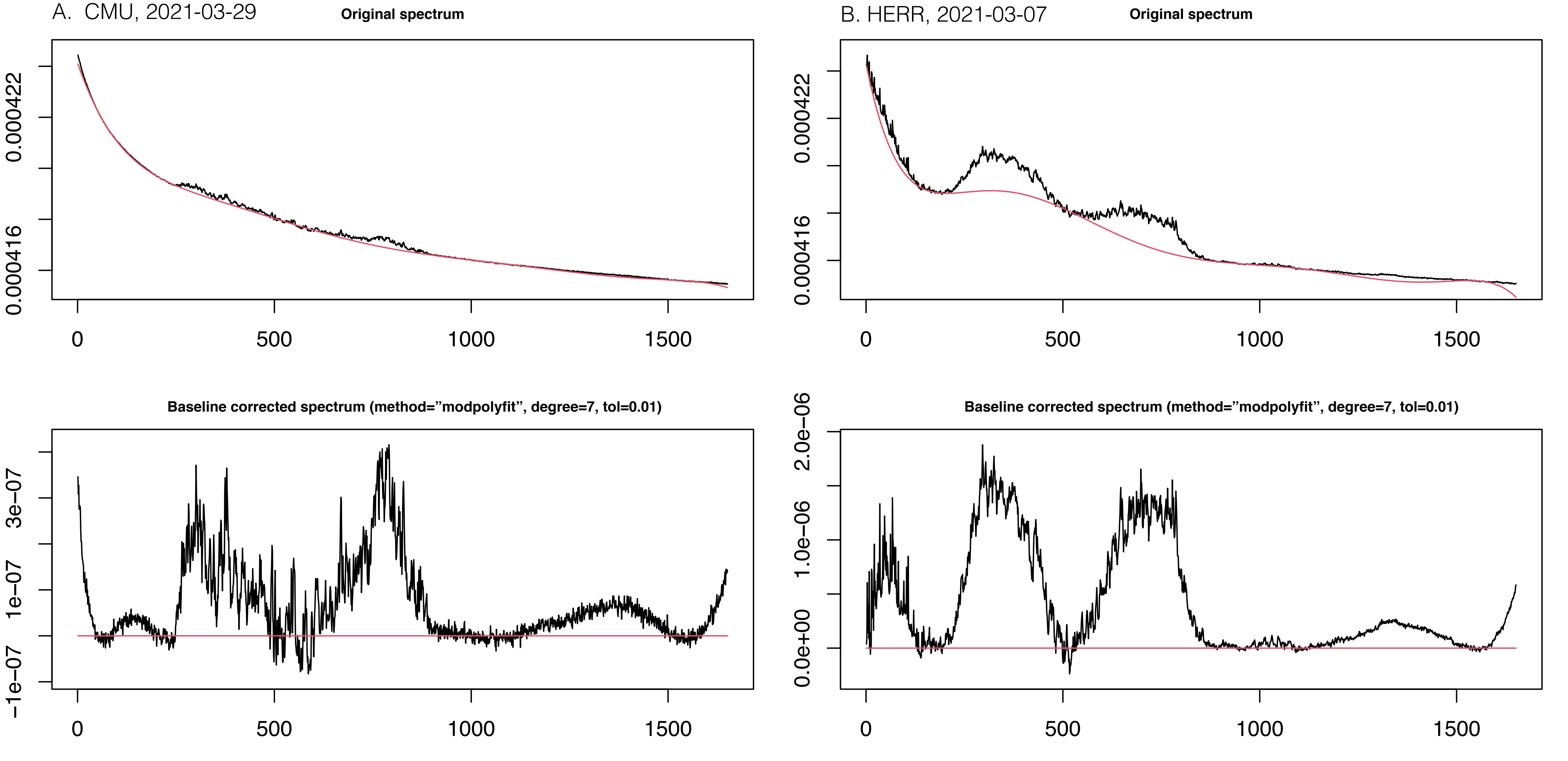

### Figure S4

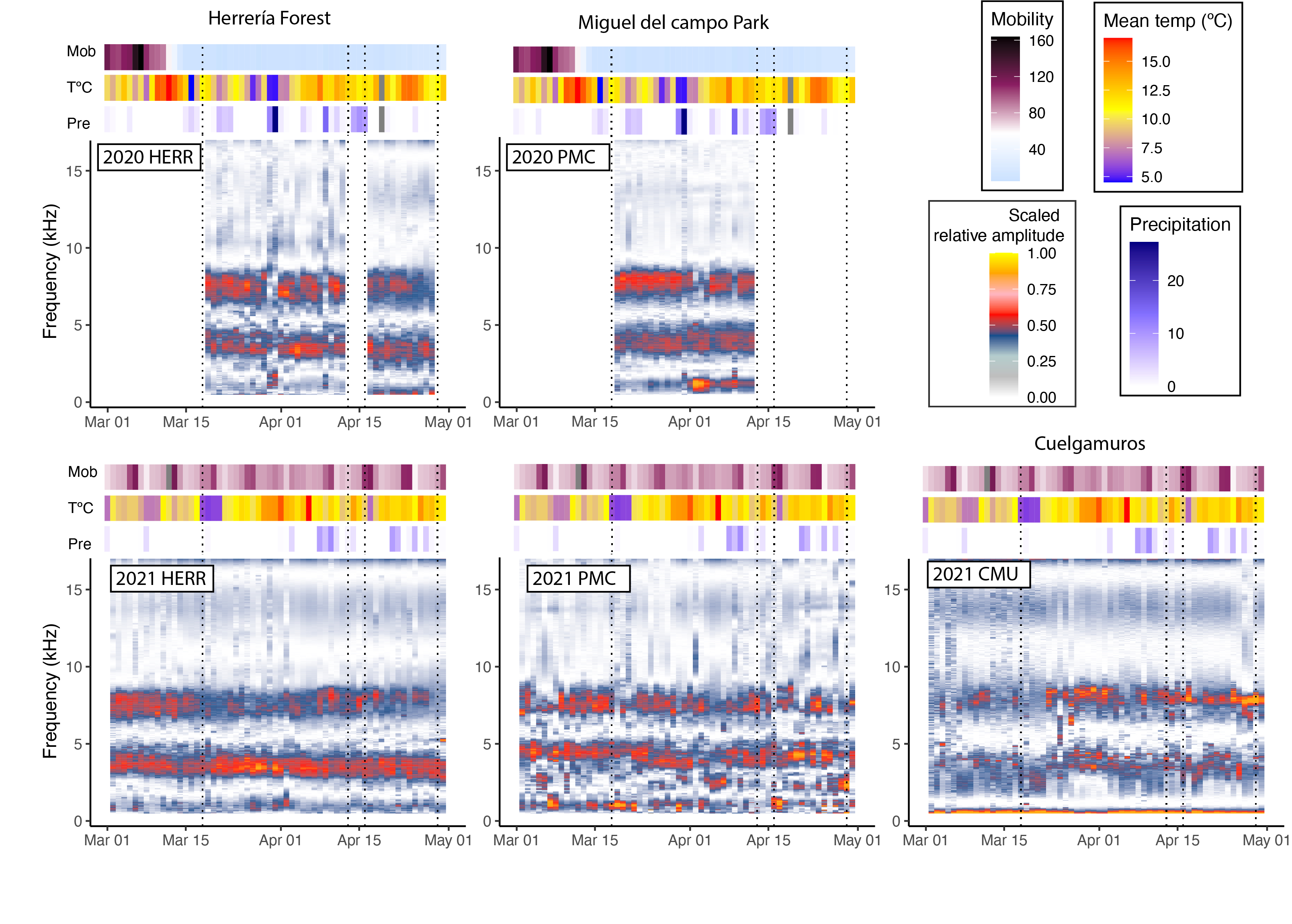

### Figure S5

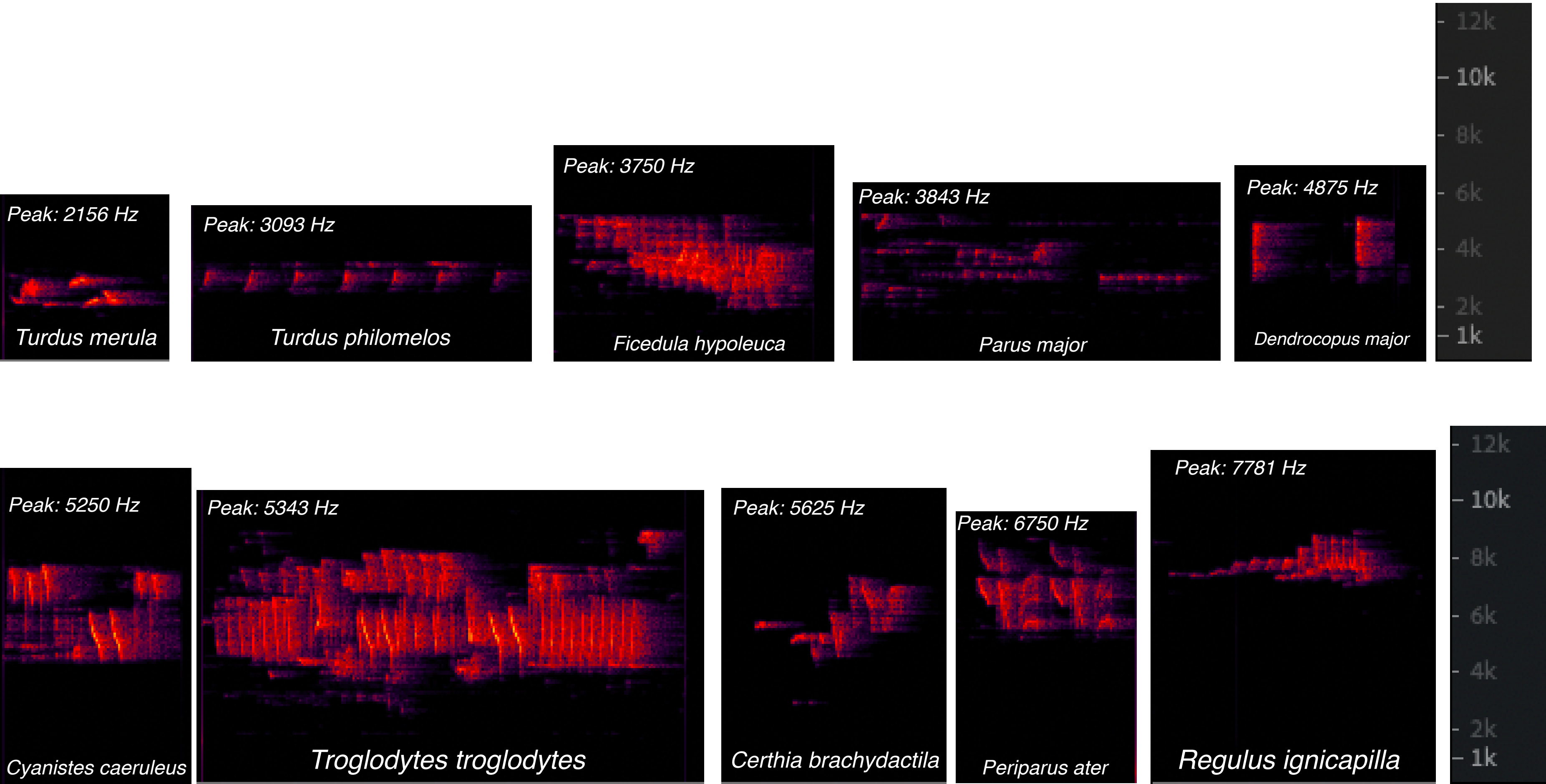

### Figure S6

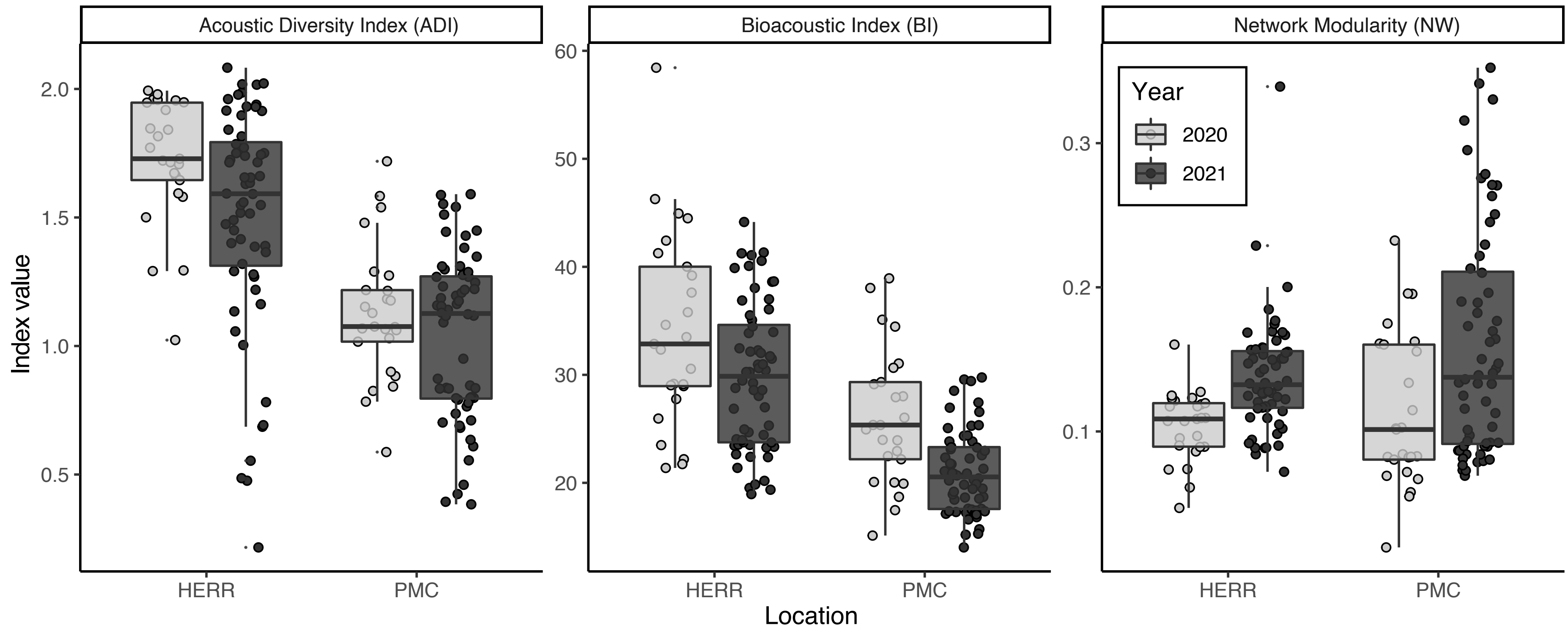
