## Supplementary Information SI1 for "A tale of two springs: contrasting forest soundscapes during the COVID-19 lockdown (2020) and after the record snowstorm Filomena (2021) from Central Spain"

**Supplementary information 1 (SI 1). Methods for the estimation of time-adjacency network metrics**

*Soundscape to networks and Network analysis*

In this article we developed a complementary method to estimate if a soundscape has different distinct parts that are organized during time (i.e., a group of organisms singing coordinately separated from a second group; or a group of biophonies separated from a geophonic event such as wind) (**Figure S3)**. To achieve this, we aimed at constructing a “temporal adjacency network” from a soundscape, and later extract network properties such as modularity (Girvan & Newman, 2006) or average path length (Csardi, 2008)~~.~~ To get the temporal adjacency network, we split the 10 minute soundscapes in 1s fragments (max. 600 segments). For each segment, the frequency spectrum was estimated using function ‘spec’ in R package seewave (Sueur, Aubin & Simonis, 2008), and rounding frequency resolution to two decimals (i.e., 4.12 kHz). We kept the spectrums between 0.5 kHz and 17 kHz, and transformed the intensity along the frequencies to relative values. To classify the spectrums, first we calculated the distance between frequency spectrums by using bray-curtis dissimilarities, second we used the dissimilarity matrix as an input to a T-SNE ordination as implemented in the R package Rtsne (Krijthe, 2015), and third we used the partitioning around medoids (pam) k-clustering, following the dissimilarity-based generalisation in Hennig and Liao (Henning, 2014). Once classified, we assigned weights to adjacent clusters (max distance, 2 seconds), and summed the weights for each pair of clusters. These weights define the network edges. Finally, we filtered out the edges with the lower summed weight (below the group mean), since these may be more spurious/random than the strongest associations. With the edge list we constructed the network using the R package iGraph, and estimated two network properties: modularity (using cluster_walktrap) and average path length. The full code for this estimation is available in Github (**https://github.com/0Rudiger/time_adjacency_networks.git**), and a graphic summary in **Figure S3**.

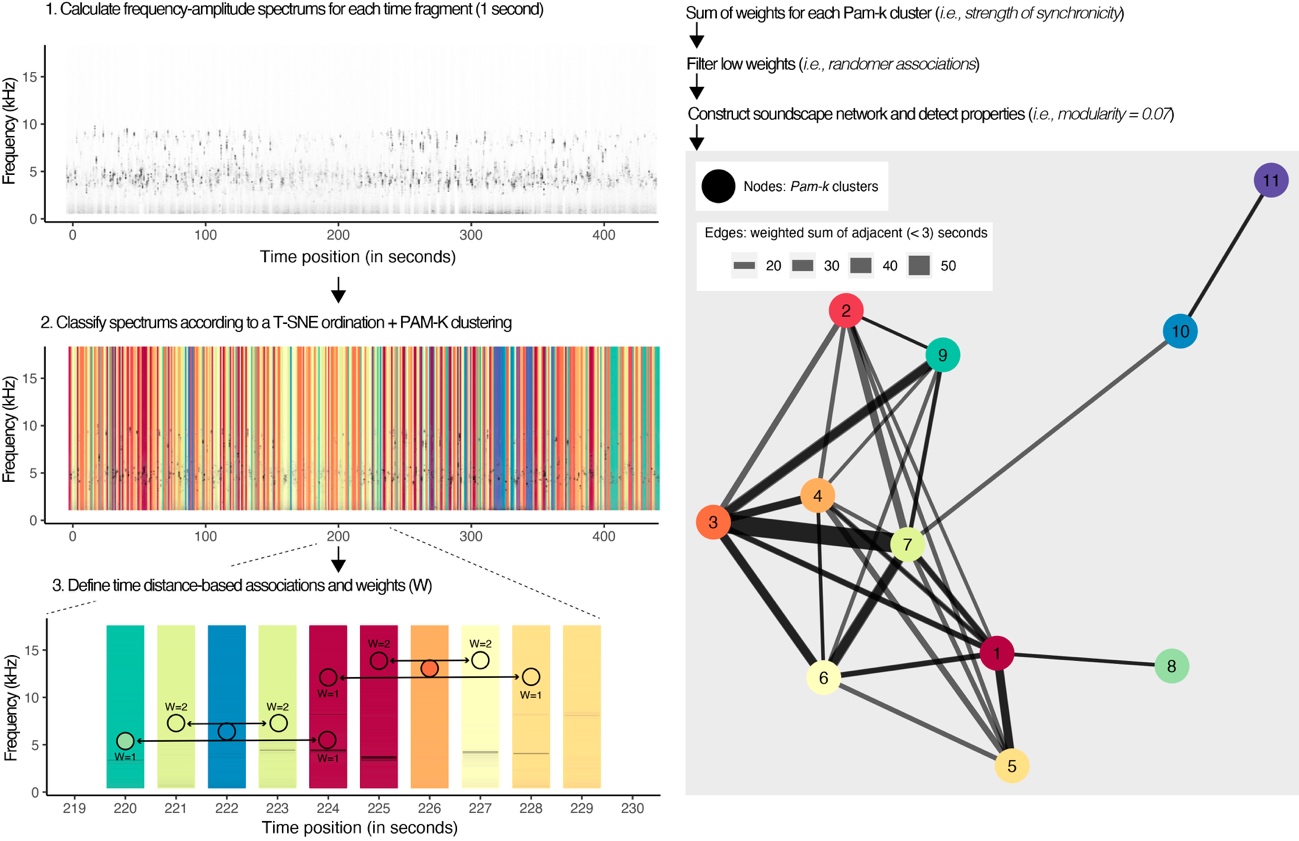

**Figure S3.** Methodological workflow for the inference of network properties from 10 minute-length soundscapes. Example figures correspond to a representative soundscape from one of the locations (HERR).
